# Lipopolysaccharide Simulations are Sensitive to Phosphate Charge and Ion Parameterization

**DOI:** 10.1101/752188

**Authors:** Amy Rice, Mary T. Rooney, Alexander I. Greenwood, Myriam L. Cotten, Jeff Wereszczynski

## Abstract

The high proportion of lipopolysaccharide (LPS) molecules in the outer membrane of Gram-negative bacteria make it a highly effective barrier to small molecules, antibiotic drugs, and other antimicrobial agents. Given this vital role in protecting bacteria from potentially hostile environments, simulations of LPS bilayers and outer membrane systems represent a critical tool for understanding the mechanisms of bacterial resistance and the development of new antibiotic compounds that circumvent these defenses. The basis of these simulations are parameterizations of LPS, which have been developed for all major molecular dynamics force fields. However, these parameterizations differ in both the protonation state of LPS as well as how LPS membranes behave in the presence of various ion species. To address these discrepancies and understand the effects of phosphate charge on bilayer properties, simulations were performed for multiple distinct LPS chemotypes with different ion parameterizations in both protonated or deprotonated lipid A states. These simulations show that bilayer properties, such as the area per lipid and inter-lipid hydrogen bonding, are highly influenced by the choice of phosphate group charges, cation type, and ion parameterization, with protonated LPS and monovalent cations with modified nonbonded parameters providing the best match to experiments. Additionally, alchemical free energy simulations were performed to determine theoretical pK*_a_* values for LPS, and subsequently validated by ^31^P solid-state NMR experiments. Results from these complementary computational and experimental studies demonstrate that the protonated state dominates at physiological pH, contrary to the deprotonated form modeled by many LPS force fields. In all, these results highlight the sensitivity of LPS simulations to phosphate charge and ion parameters, while offering recommendations for how existing models should be updated for consistency between force fields as well as to best match experiments.

## Introduction

Gram-negative bacterial infections are a significant public health threat,^1^ and are typically more difficult to treat than Gram-positive infections^2–4^ due largely to the presence of a second, outer membrane surrounding their peptidoglycan cell wall and plasma membrane. This outer membrane is highly asymmetric, containing predominately phospholipids in the inner leaflet, while the outer leaflet is rich in lipopolysaccharides (LPS).^5, 6^ LPS are structurally dissimilar from glycerophospholipids, and are composed of three regions: lipid A, which contains multiple saturated hydrocarbon chains and acts as the hydrophobic anchor; the core region, a collection of branched oligosaccharides that are often phosphorylated; and the O-antigen, a polymer of repeating saccharide units. The large number of anionic groups present in LPS imparts a net negative charge to the molecule, and leaflets are stabilized by a network of divalent cations bridging these moieties. ^7, 8^

LPS modification processes, such as the PhoPQ system depicted in Figure 1, reduce this charge through adornment of the lipid A phosphate groups. ^9–11^ In *S. enterica*, these modifications are activated by a variety of environmental stimuli, such as a low concentration of divalent cations, ^12, 13^ acidic conditions, ^14, 15^ hyperosmotic stress, ^16^ or antimicrobial peptide presence, ^17, 18^ indicating that modifications may confer a survival advantage in such conditions. Additionally, previous simulations have shown that the presence of aminoarabinose disrupts the cation network, allowing direct inter-lipid hydrogen bonding to instead stabilize the leaflet and potentially reducing the reliance on divalent cations for stability.^19^

**Figure 1:**
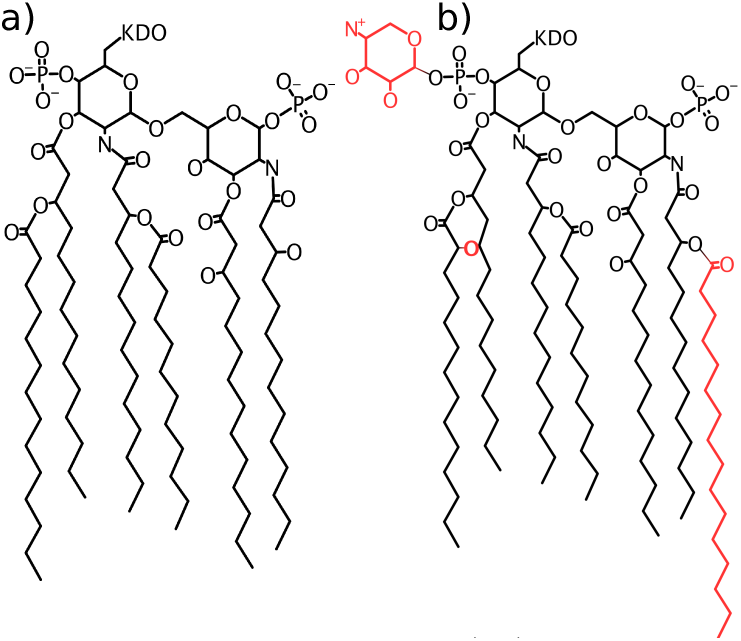
Structure of LPS used in this study. (A) Structure of unmodified lipid A. (B) Modification by the *PhoPQ* regulatory system results in three key additions to the lipid A structure, shown in red. Throughout the text, we refer to LPS containing these lipid A modifications as mLPS to distinguish it from the unmodified chemotypes.

Experimentally, LPS monolayers and bilayers display significant structural changes upon monovalent or divalent cation inclusion. X-ray reflectivity experiments on *Salmonella* LPS exhibited a clear trend of larger lamellar repeat periods with Ca^2+^ and Mg^2+^ compared to Na^+^, regardless of chemotype. ^20^ X-ray diffraction on monolayers of similar LPS species showed a much lower compressibility for monolayers with Ca^2+^ than those with Na^+^.^21^ Furthermore, monolayers of *E. coli* LPS revealed significantly smaller lipid areas in the presence of 20 mM Ca^2+^ with 100 mM NaCl, compared to 100 mM NaCl alone or no ions, ^22^ indicating that the type of ion species has a larger influence on the monolayer structure than the ionic strength alone. The neutron scattering density profiles of Kucerka *et al.* revealed decreased water penetration into the LPS core in Ca^2+^-containing LPS bilayers only, compared to those containing either Na^+^ or Mg^2+^, despite the similar increased tail ordering observed with either divalent cation. ^7^ This intriguing difference between Ca^2+^ and Mg^2+^ may be attributable to calcium’s lower hydration energy than magnesium, meaning that less energy is required to remove its hydration shell. In all, these results demonstrate a clear condensing effect of divalent cations, such as Ca^2+^ and Mg^2+^, compared to Na^+^. This condensing effect is manifested through smaller lipid areas, decreased tail mobility, and increased molecular packing.

LPS models have been parameterized for all major families of molecular dynamics (MD) force fields, ^23–28^ and outer membrane simulations with LPS are increasingly more common. ^29–33^ However, the performance of common ion force fields with these LPS models has rarely been evaluated, and inconsistencies exist between different parameterizations. The GROMOS-based force field of Pontes *et al.*, when simulated with Ca^2+^ or Na^+^, was found to only form a stable lamellar phase in the presence of Ca^2+^; ^24^ simulations with Na^+^ resulted in a clear transition from lamellar to nonlamellar structures. Kim *et al.*, utilizing the CHARMM LPS force field, ^25^ reported a compaction upon inclusion of K^+^ or Na^+^ compared to Ca^2+^ for all five LPS chemotypes simulated,^34^ contrary to experiment. The closest match to experiment comes from the GLYCAM LPS^23^ force field– simulations of asymmetric LPS bilayers displayed a decreased lipid area with divalent cations compared to monovalent cations; ^8^ however, the bilayer was not stable in the presence of K^+^, breaking down within the first 50 ns of the simulation.

The troubling inconsistencies between these force fields may arise in part from the different ion and water models used. However, additional discrepancies exist between LPS parameterizations– there is no consensus on the charge state of phosphate groups within LPS and different force fields assign different net charges. The atomistic GLYCAM,^23^ GROMOS-based, ^24^ and AMBER^27^ force fields, as well as MARTINI-compatible coarse grained (CG) models built off them, ^35, 36^ assign a charge state of −1 to the lipid A phosphate groups. In contrast to this, the CHARMM LPS force field^25, 34^ and corresponding CG models^26^ treat the lipid A phosphate groups as fully deprotonated, with a charge of −2 per phosphate. While no titration data exist for lamellar LPS, solubility experiments on the Re chemotyle of *E. coli* LPS determined the pK*_a_* values in solution as 8.6 for the first lipid A deprotonation and 10.8 for the second deprotonation,^37^ indicating that the fully protonated state (charge of −1 per phosphate group and −4 per LPS) should dominate in solution at physiological pH.

In this study, we investigate the effects of both ion type and phosphate charge on four distinct chemotypes of *S. enterica* LPS (Figure 2) to better characterize these force field discrepancies as well as to study the protective effects that LPS modifications confer. We report that simulation results, such as the area per lipid, core hydration, and inter-lipid hydrogen bonding are highly influenced by the choice of protonation state, cation type, and ion parameterization. Complimentary alchemical free energy simulations and ^31^P-NMR spectroscopy experiments determine that the protonated state (−1 per phosphate group) dominates at or near physiological pH. As hypothesized, simulations with modified LPS show a decreased sensitivity to ion substitutions when compared to unmodified LPS forms. Overall, these results demonstrate the sensitivity of LPS simulations to parameterization differences and offer guidelines for how existing LPS parameterizations should be modified for better agreement with one another and experiment.

**Figure 2:**
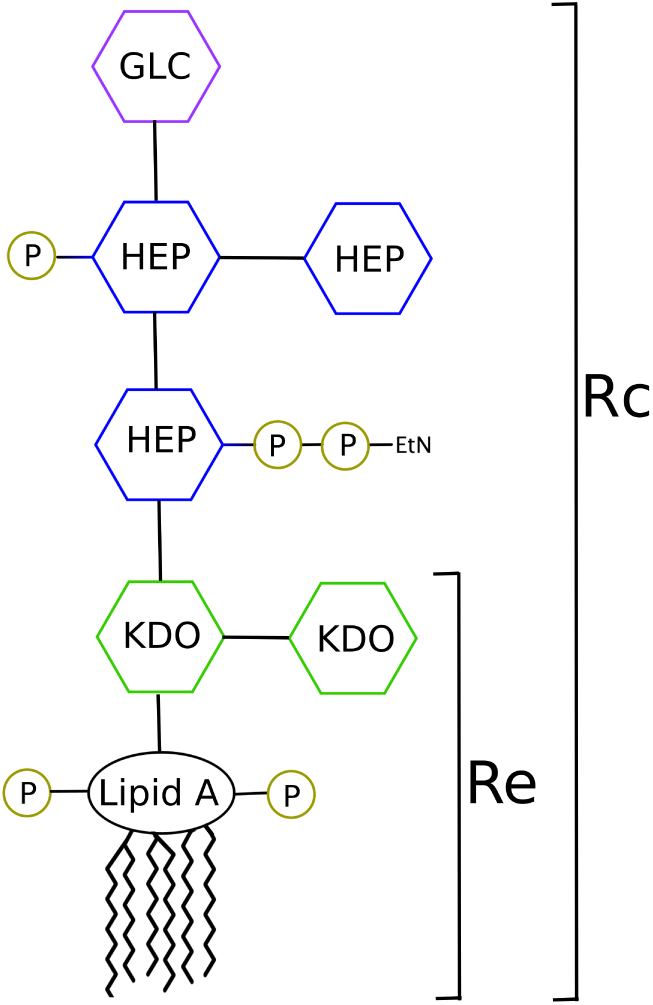
Schematic of the *S. enterica* LPS Rc core. The Re chemotype contains only the two KDO sugars. Abbreviations: KDO, 2-keto-3-deoxyoctulosonic acid; Hep, heptose; Glc, glucose; EtN, ethanolamine; P, phosphate.

## Methods

### System preparation

4×4 Rc LPS bilayers were constructed by removing 20 lipids per leaflet from a pre-equilibrated 6×6 Rc LPS bilayer; this 4×4 system was then solvated and equilibrated for 2.0 µs. A 4×4 Rc modified LPS (mLPS) system was constructed and equilibrated in a similar manner. No significant differences were observed between these 4×4 LPS simulations and similar 6×6 systems (Section S2 in the Supporting Information). These equilibrated 4×4 Rc systems were stripped of all water and ions, then resolvated with neutralizing counterions added to the bulk solvent. In all cases, enough water was added to ensure a *∼*1.5 M monovalent cation concentration, and the same solvent box was used for the simulations with divalent ions.We note that LPS are highly charged molecules (−9 to −4 net charge depending on the chemotype), necessitating a large number of counterions to maintain a neutral simulation box. While the initial ion concentration in bulk solvent upon system construction is quite high, these cations quickly move to saturate the LPS core within the first few nanoseconds of the solvent equilibration described below. The number of cations in the bulk solvent during the production portion of simulations is minimal for all systems studied (Figure S6).

Simulations utilized either the default CHARMM ion parameters of Beglov and Roux, ^38^ the NBFIX calcium parameters of Roux and Rong as reported by Kim *et al.*,^34^ or the CUFIX parameters of Yoo and Aksimentiev^39^ for all neutralizing counterions. All systems utilized the LPS parameter set of Wu *et al.* ^25^ with modifications treated as described previously,^19^ the C36 force fields for lipids, ^40, 41^ modified Lennard-Jones parameters for sodium ion interactions with certain lipid oxygens, ^42^ and TIP3P water. ^43^ Lipid A phosphate groups with a charge of −1 were parameterized by analogy to the C36 general force field, ^44^ using methylphosphate as a template; the parameters used are given in Table S1. We note that in all Rc systems, phosphate groups in the core oligosaccharide region retained their default charge of −2, regardless of the treatment of the lipid A phosphate group charge; Re systems do not contain any core phosphate groups. Systems were converted to AMBER-compatible format using chamber ^45^ in ParmEd, then minimized, heated, and equilibrated for 5 ns with LPS sugars restrained to allow the water density to equilibrate. This 5 ns solvent equlibration was sufficiently long to allow hydration of the LPS core; furthermore, in all simulations the majority of counterions were associated with the LPS core by the end of this solvent equilibration.

### Conventional molecular dynamics simulations

Initially, 56 different systems were simulated, accounting for all possible combinations of four LPS chemotypes, seven ion models, and two different phosphate net charges (Table 1). Four additional simulations were performed of Re LPS with −1 phosphate charges interacting with an excess of Ca^2+^, K^+^, Mg^2+^, or Na^+^ to verify the results of these lower charge simulations were not a consequence of having fewer cations present (Section S1). All conventional MD (cMD) production simulations were performed with the GPU-accelerated version of *pmemd* in AMBER 18, ^46^ and were carried out for 3.0 µs per system. System temperatures were controlled at 310 K using Langevin dynamics, with a collision frequency of 1.0 ps*^−^*^1^. The pressure was maintained at 1.0 bar by means of semi-isotropic coordinate scaling, with z decoupled from the xy dimensions, and utilized the Berendsen barostat^47^ with a relaxation time of 1.0 ps. All hydrogen bonds were constrained using SHAKE.^48^ A 12.0 Å cutoff was used, with interactions smoothly switched to zero over 10–12 Å. Long-range electrostatics were treated using the particle mesh Ewald (PME) method ^49^ with a grid spacing of 1.0 Å.

**Table 1:** Enumeration of the different system options used for cMD simulations. * Refer to Figures 1 and 2 for chemotype depictions. †Default CHARMM parameters of Beglov and Roux. ^38^ ‡NBFIX calcium parameters of Roux and Rong as reported by Kim *et al.* ^34^ CUFIX parameters of Yoo and Aksimentiev.^39^ The exact Lennard-Jones parameters used are given in Table S6.

| System Feature | Number | Specification |
| --- | --- | --- |
| LPS chemotype* | 4 | Rc LPS, Rc mLPS, Re LPS, Re mLPS |
| Ion model | 7 | Ca<sup>2+</sup>, K<sup>+</sup>, Mg<sup>2+</sup>, Na<sup>+</sup> †<br>NBFIX Ca<sup>2+</sup> ‡<br>CUFIX K<sup>+</sup>, CUFIX Na<sup>+</sup> ° |
| Lipid A phosphate charge | 2 | -2, -1 |

### Alchemical free energy simulations

Alchemical free energy simulations were performed in NAMD 2.13, ^50^ utilizing interleaved double-wide sampling to allow sampling in both the forward and reverse direction at each lambda window. Simulations were performed at fifteen windows with λ= 0.0, 0.01, 0.05, 0.1, 0.2, …, 0.9, 0.95, 0.99, 1.0. Here, λ= 0.0 corresponds to the state where the proton in question is present, while λ= 1.0 corresponds to the deprotonated state. Four distinct alchemical transformations were performed: from the −4 to −5 charge state by deprotonation of either the P*_A_* or P*_B_* phosphate of Re LPS, and from the −5 to −6 charge state by deprotonation of the second phosphate group (Figure 3). Each of these four alchemical transformations were performed in three different environments: solvent, a fully deprotonated (−6 state) Re LPS bilayer, and a doubly protonated (−4 state) Re LPS bilayer.

**Figure 3:**
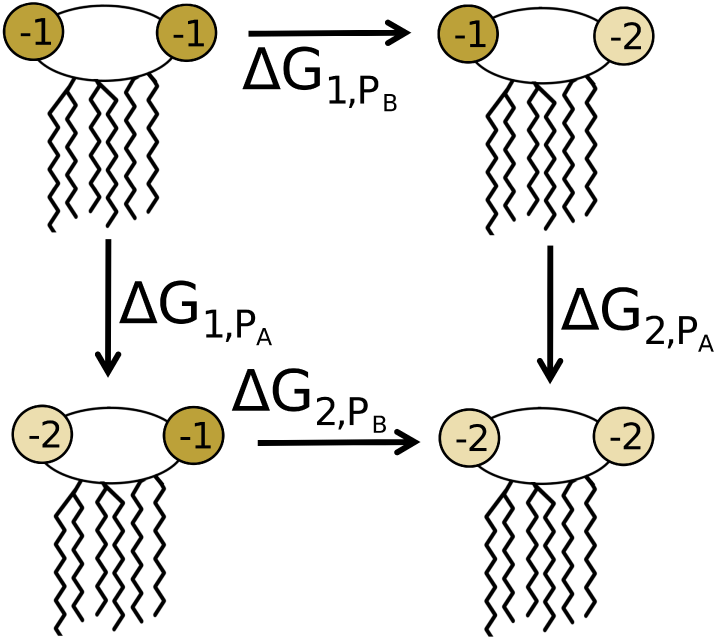
Depiction of the four alchemical transformation pathways explored using free energy simulations. These correspond to deprotonation of either the P*_A_* or P*_B_* phosphate, with the other phosphate group either protonated or deprotonated.

Each lambda window was equilibrated for 1 ns, followed by 10 ns of production MD. Soft-core vdW potentials and a delayed introduction of electrostatics (lambda *>* 0.1) were performed to avoid infinite electrostatic or vdW interactions at end points. The bilayer simulations were performed five times each for robust error analysis, while the solvent simulations were repeated three times.

### Simulation analysis

Trajectory analysis was performed over the final 1.0 µs of each cMD simulation, to allow ample time for bilayer equilibration. Lipid area, hydrogen bonds, carbon-deuterium order parameters, and electron density profiles along the bilayer normal were calculated using *CPPTRAJ* ^51^ from AmberTools 17.^52^ Carbon-deuterium order parameters are reported as *|S*_CD_*|*. All hydrogen bond calculations utilized a distance cutoff of 3.0 Å and an angle cutoff of 135°. Cation coordinating groups were determined using a distance-based cutoff, calculated in VMD^53^ and updated every 20 frames (200 ps). The cutoff used varied depending on the ion type and corresponded to the distance of the first minima in the ions’ radial distribution function (RDF) – 3.0 Å for calcium and sodium, 3.5 Å for potassium, and 2.5 Å for magnesium. Cation radial distribution functions (RDFs) were calculated using LOOS,^54^ and coordination numbers were determined by integration of the resultant RDF through the first peak. All errors are reported as standard error of the mean.

The Bennett acceptance ratio (BAR)^55^ free energy estimator in the ParseFEP module of VMD^56^ was used to analyze all alchemical simulations. The full thermodynamic cycle utilized for these calculations is depicted in Figures 3 and 4. Here, the ΔG of each deprotonation, either in water (ΔG*_wat_*) or in the bilayer (ΔG*_bil_*), is calculated from the free energy simulations with removal of a single proton. The change in energy between deprotonation in the bilayer environment and deprotonation in water is given by ΔΔG = ΔG*_bil_* − ΔG*_wat_*. From this, the pK*_a_* shift that arises as a result of this change to the phosphate group’s environment can be directly calculated:

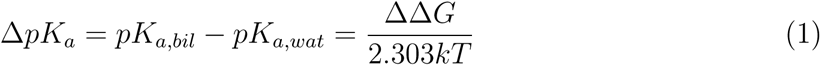

**Figure 4:**
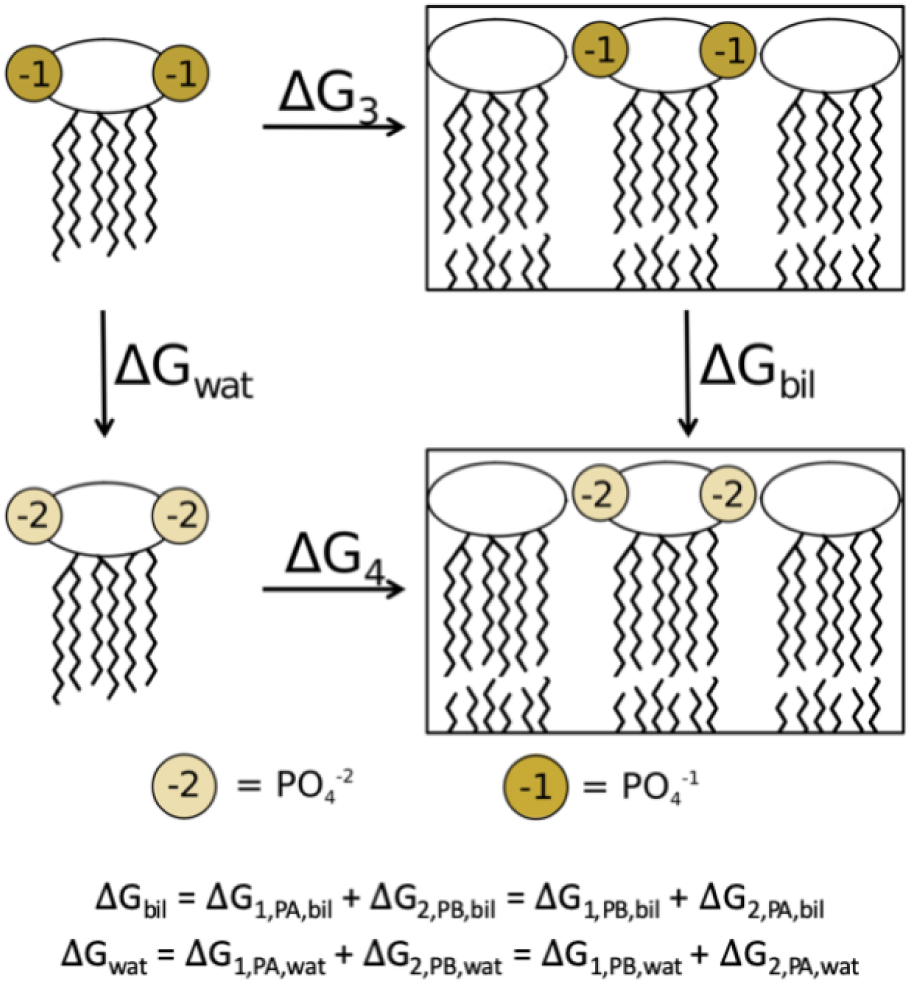
Thermodynamic cycle utilized in the alchemical simulations. Δ*G_wat_* and Δ*G_bil_* are calculated as the sum of the individual deprotonation steps shown in Figure 3.

where *k* is the Boltzmann constant and T is the system temperature. Since the pK*_a_* for Re LPS phosphate groups in water is known, this method allows estimation of the pK*_a_* in the bilayer.

### Experimental pH studies of LPS using solid-state NMR

Pure LPS has been observed to form both unilamellar and multilamellar vesicles, ^57, 58^ making it suitable for ^31^P NMR experiments. LPS *E. coli* strain R515, corresponding to the Re chemotype, was purchased from AdipoGen Life Sciences (Liestal, Switzerland) in 1 mL vials of sterile aqueous solution (1 mg/mL). These solutions were lyophilized, rehydrated with nanopure water, and consolidated to get 5 samples with LPS mass of 2 mg each. The samples were then lyophilized again before resuspending to a LPS concentration of 20 mM in 57 mM Michaelis barbital sodium-acetate buffer ^59^ at pH 1.99, 5.02, 6.99, 8.98, and 10.46. Solid CaCl_2_*·*2H_2_O was added to reach a Ca^2+^ concentration of 50 mM. Each solution was incubated at 40°C for 30 minutes before packing the samples into 2.5 mm rotors (Bruker, Billirica, MA) or 5 mm glass tubes (New Era, Vineland, NJ) for NMR analysis.

All NMR experiments were performed on a 17.6 T (750 MHz) wide bore (Bruker, Billerica, MA) spectrometer with a variable temperature set to 32°C with a Bruker BVT-3000 temperature controller. ^31^P experiments employed SPINAL-64 ^1^H decoupling^60^ during acquisition, with a nutation frequency of 36 kHz (MAS experiments) or 78 kHz (static experiments). Static experiments were performed using a low-electrical field probe (Black Fox, Inc., Tallahassee, FL) and MAS experiments were carried out using a Bruker BL2.5 HX 2.5 mm MAS probe. In MAS experiments, spinning was regulated at 15,000 *±* 10 Hz using a Bruker MAS II pneumatic MAS controller. The recycle delay was 2 s (MAS experiments) or 3 s (static experiments) and the 90°-pulse was 2.5 µs (MAS experiments) or 6 µs (static experiments). MAS spectra were indirectly referenced to adamantane externally, assuming the downfield peak of 38.48 ppm.^61^ NMR spectra were processed with 500 Hz (MAS experiments) or 1000 Hz (static experiments) exponential apodization and zero filled and left shifted prior to Fourier transformation. Spectra were then subjected to polynomial baseline subtraction. All samples exhibited signals consistent with stable lamellar structures (Figure S3), and displayed the same characteristic lineshape as that of phospholipids in the lamellar phase.^62^

## Results

Average bilayer properties for all systems are given in Tables S2 and S3 in the Supporting Information. In all the measures studied, the bilayer structure and organization was greatly affected by the cation type included and the charge on the lipid A phosphate groups.

### Lower phosphate charges lead to bilayer compaction

Protonation of the lipid A phosphate groups resulted in a markedly more compact, ordered bilayer, regardless of the LPS chemotype or cation type included. Simulations with 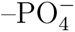 led to a lower area per lipid and a thicker hydrophobic region when compared to equivalent simulations with 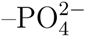 (Figure 5). For example, in the case of Re LPS the average area per lipid decreased from 164 *±* 0.4 Å^2^ to 148 *±* 0.2 Å^2^ with a concomitant thickening of the leaflet hydrophobic thickness from 12.8 *±* 0.1 to 13.6 *±* 0.1 Å. Additionally, tail order parameters demonstrated a distinct ordering in LPS simulations when 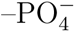 was implemented in place of the standard CHARMM 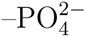 (Figure S4); this difference was less pronounced in mLPS systems, likely due to the already increased ordering that palmitoylation confers. ^19^ Finally, increased inter-lipid hydrogen bonding was observed in LPS systems with the reduced phosphate charge, while the change in mLPS systems was not statistically significant for this metric.

**Figure 5:**
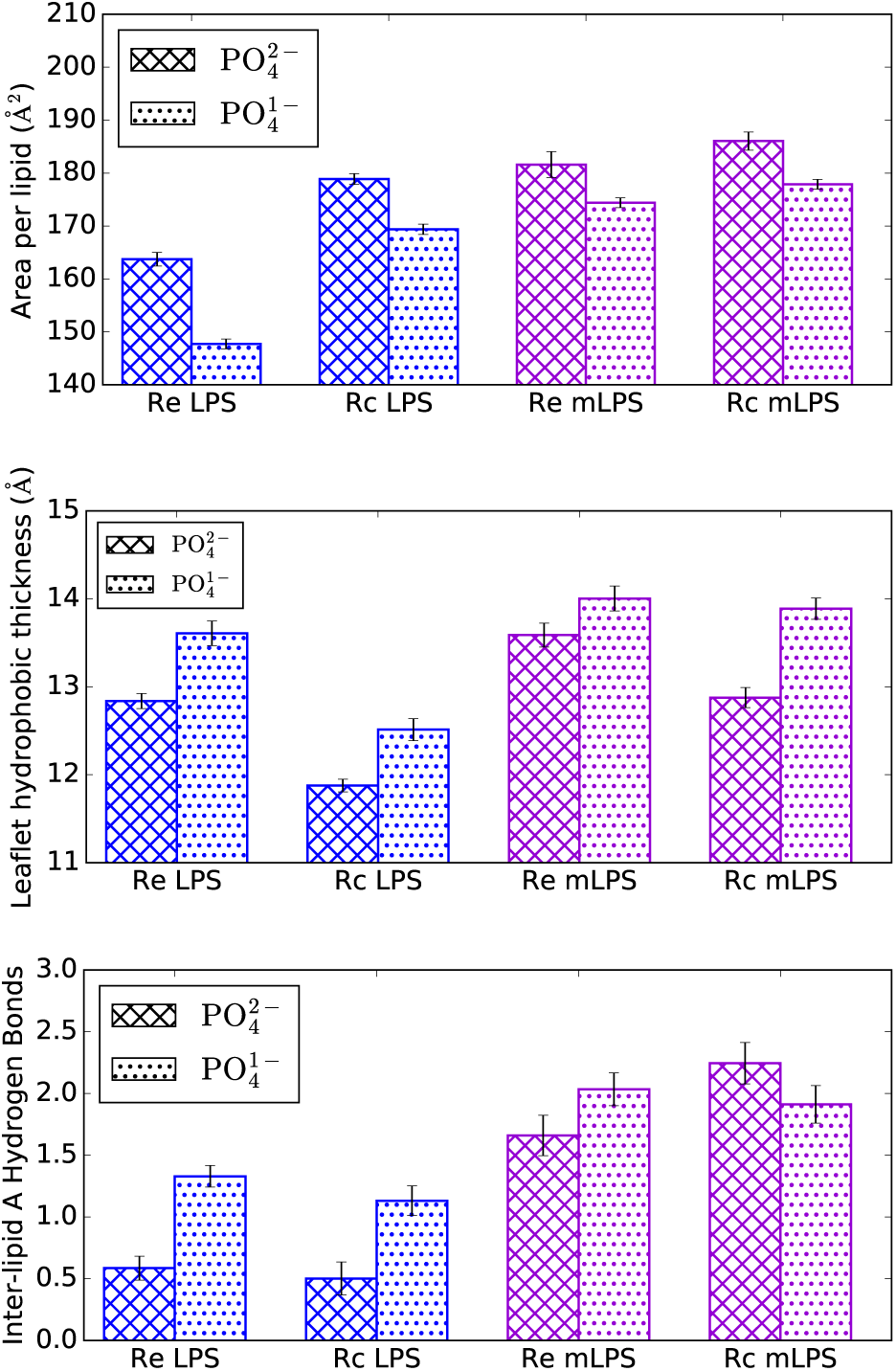
Effect of lipid A phosphate charge on LPS area (top), per-leaflet hydrophobic thickness (middle), and inter-lipid A hydrogen bonding (bottom). Data are shown from the simulations with Ca^2+^.

Furthermore, simulations with reduced lipid A phosphate charges resulted in a less hydrated LPS core, especially in the region of those phosphate groups (Figure S5 and Table S4); it is unclear whether this is an effect from the decreased lipid area or a result of fewer hydrated cations present in the core. However, regardless of the phosphate charge or core hydration, counterions remained strongly associated with the LPS core and were rarely present in the bulk solvent (Figure S6). Overall, these results demonstrate that differing lipid A phosphate protonation states lead to clear structural variations in the simulation outcome.

### Cation type greatly affects LPS bilayer properties

Lipid packing was surprisingly highly affected by the cation type included in the simulations. The standard CHARMM ion parameters, which include Na^+^ NBFIX terms, resulted in simulations with Mg^2+^ counterions typically having the largest area per lipid, while simulations with Na^+^ tended to form the most compact bilayer (Figure 6). This trend was roughly the same regardless of LPS type or phosphate charge. Coupled with these area per lipid changes were concomitant changes to the leaflet hydrophobic thickness and inter-lipid A hydrogen bonding, with lower lipid areas corresponding to more ordered lipid tails.

**Figure 6:**
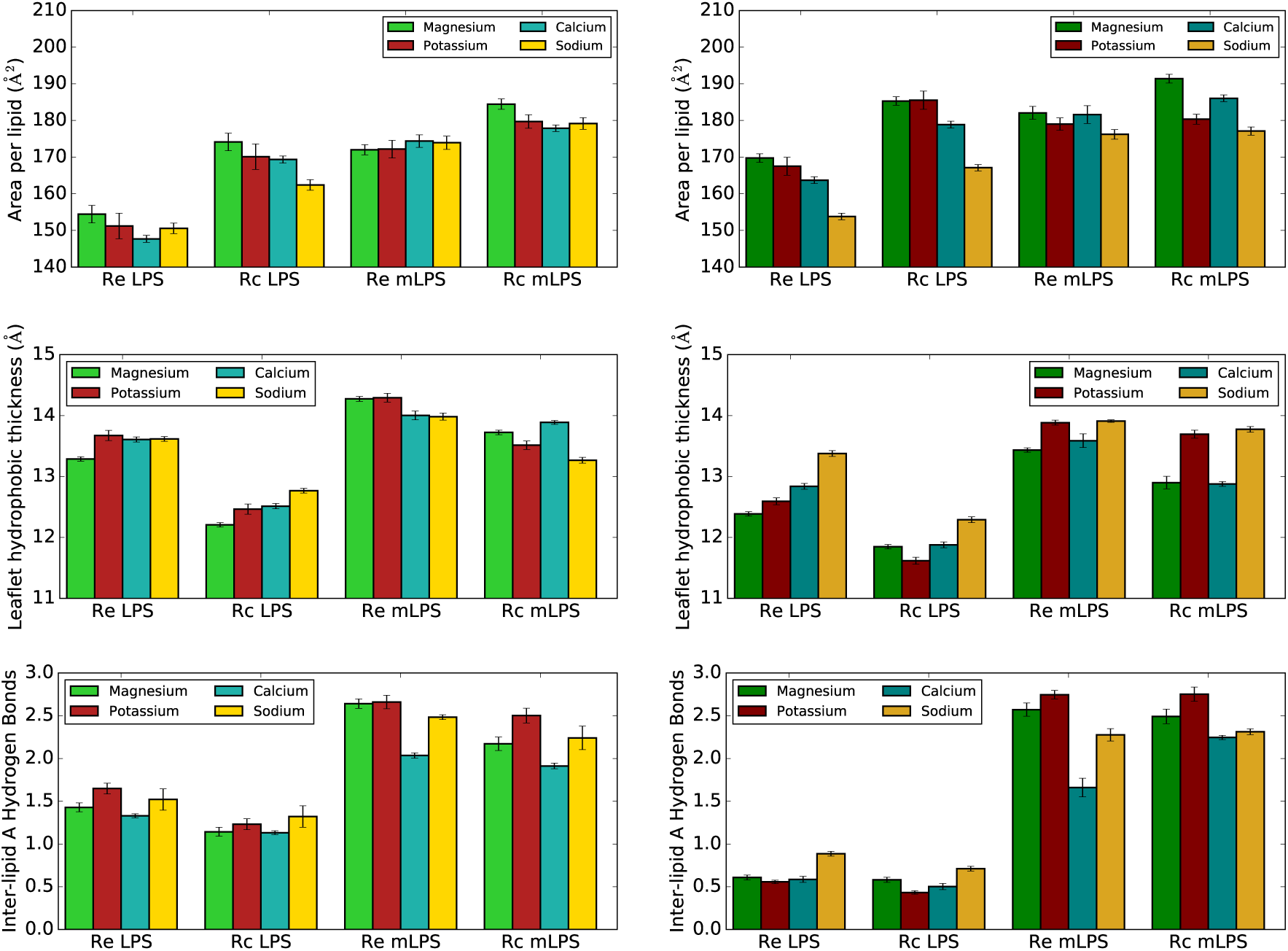
Effect of cation inclusion on LPS area (top), per-leaflet hydrophobic thickness (middle), and inter-lipid A hydrogen bonding (bottom). Systems with 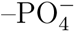 are shown on the left, while systems with 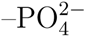 are on the right.

Utilization of the Ca^2+^ NBFIX^34^ and Na^+^/K^+^ CUFIX^39^ parameter sets led to significantly better agreement with experimental lipid area trends. Simulations with these parameter sets, which refine cation interactions with phosphate and carboxyl oxygen atoms, correctly recover the experimentally-observed increased lipid area upon monovalent cation inclusion (Figure 7). This lipid area trend reversal is less striking in mLPS simulations; however, no experimental data exist for these LPS types, and the presence of aminoarabinose may likely alter the expected area trend. These results indicate that the Na^+^/K^+^ CUFIX ion parameter sets may be the most appropriate for use with LPS simulations. Additionally, the observed changes unequivocally demonstrate that small modifications to specific non-bonded parameter pairs is sufficient to profoundly affect bilayer structural properties, indicating that further refinement to experiments may be possible.

**Figure 7:**
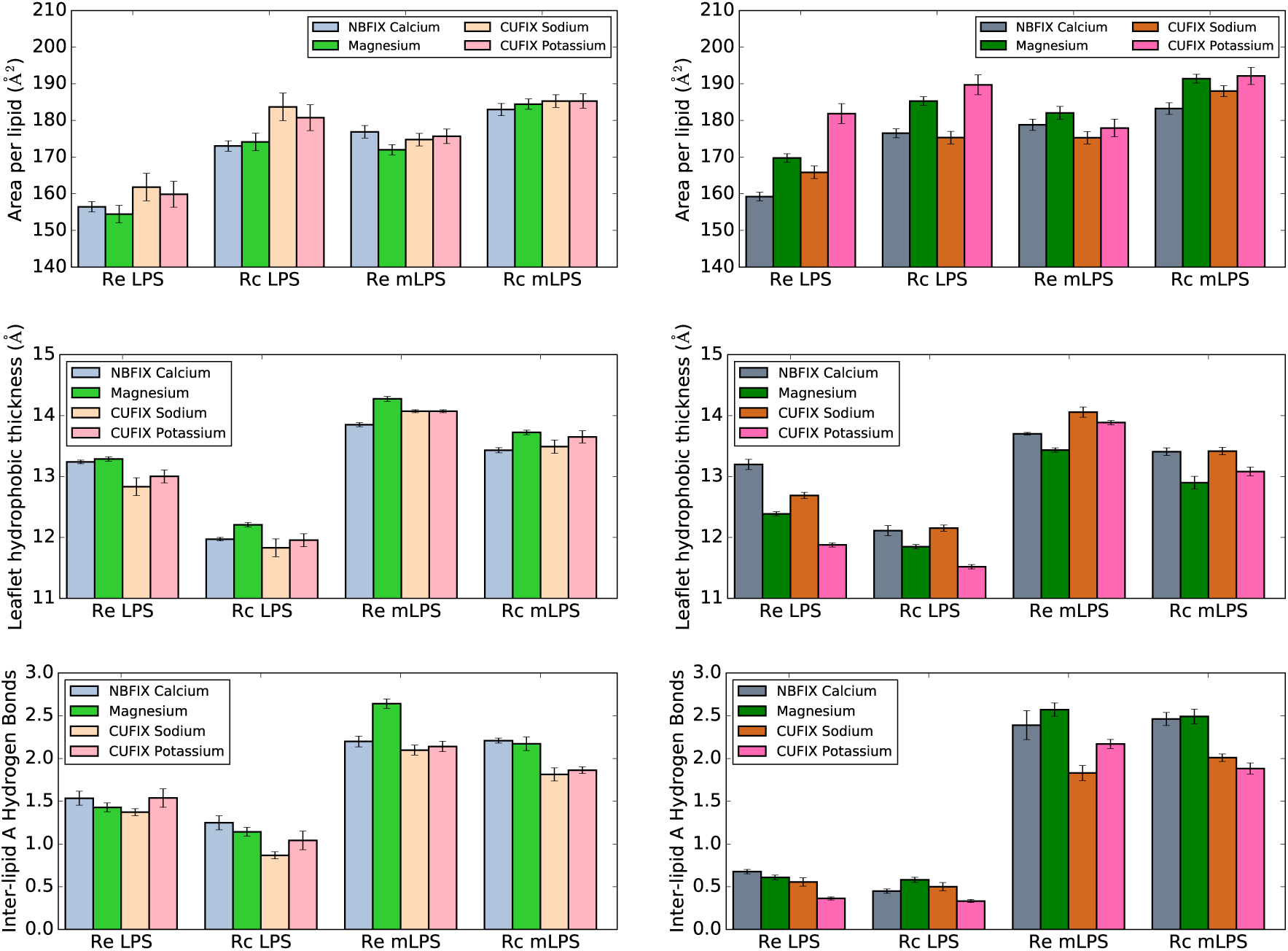
Effect of NBFIX/CUFIX cation inclusion on LPS area (top), per-leaflet hydrophobic thickness (middle), and inter-lipid A hydrogen bonding (bottom). Systems with 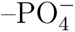 are shown on the left, while systems with 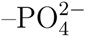 are on the right.

### Modification confers resilience to ion-induced membrane changes

Regardless of the chemotype or phosphate charge parameterization, simulations with PhoPQ-mediated modifications present exhibit more stable per lipid areas than those without modifications present (Figure 8). For example, in Re LPS with 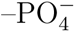, the percent change to the lipid area is 5.4% upon inclusion of different cations compared to Ca^2+^, while the percent change is only 0.9% in Re mLPS. This trend holds for both Rc and Re chemotypes with either phosphate charge. Interestingly, the percent area change is roughly the same for each chemotype when the 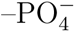 and 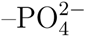 cases are compared. Overall, these results indicate that, as hypothesized previously,^19^ lipid A modifications help stabilize the bilayer structure in the absence of divalent cations and may reduce reliance on these cations for stability.

**Figure 8:**
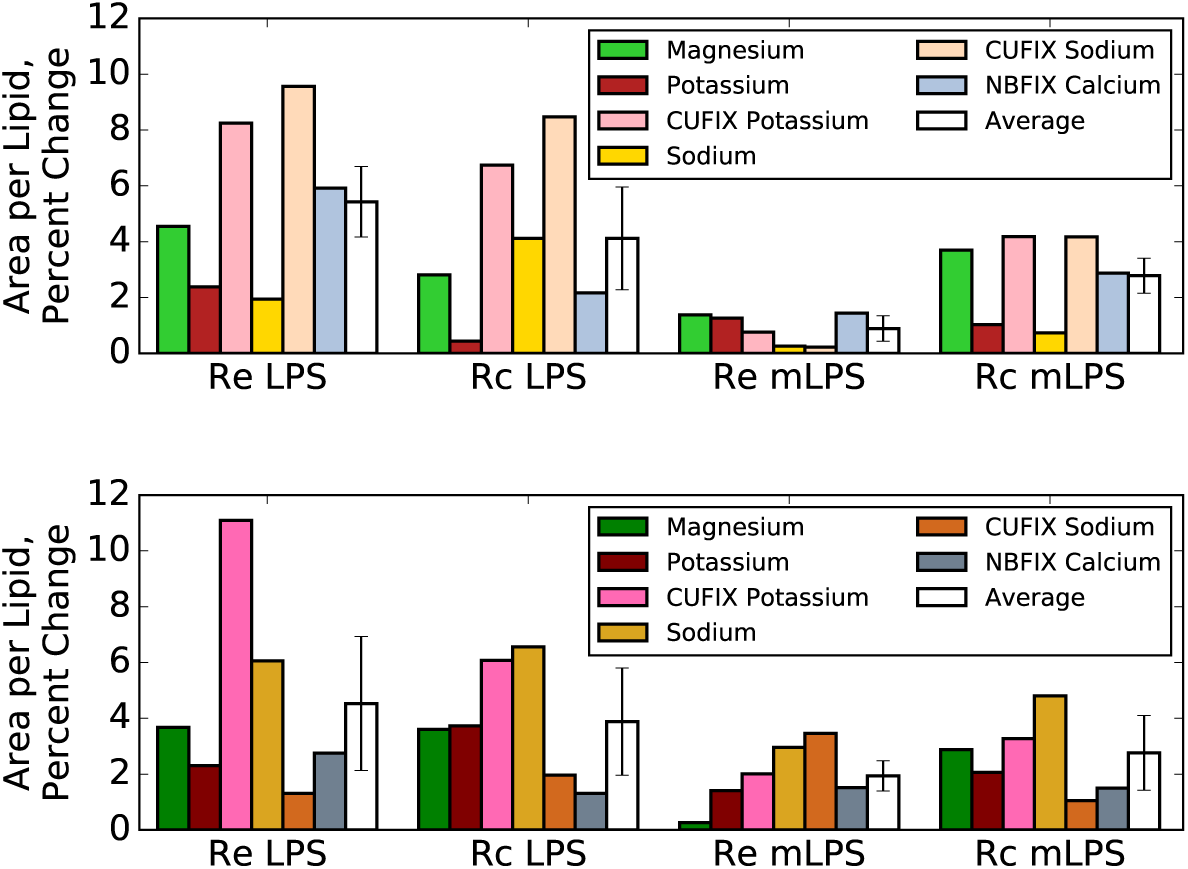
Percent change in area per lipid different ion species for all four chemotypes studied, compared to simulations with Ca^2+^ present. Systems with 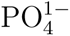 are shown in the top panel, while systems with 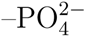 are on the bottom.

### Bilayer presence shifts lipid A phosphate pK*_a_* values

Free energy differences for each of the four possible deprotonation steps studied (Figure 3) are given in Table S5. For each step, there was good agreement between all three or five calculation replicates performed, as indicated by the small standard errors of the mean. Additionally, energy estimates for each lambda window were well converged on the simulation timescales used here, and there were no deviations between forward and backward free energy estimates. From these ΔG results, the ΔΔG between deprotonation in the bilayer and deprotonation in solution can be calculated. These ΔΔG values for the first (−4 to −5 charge state) and second (−5 to −6) deprotonations are given in Table 2, along with the corresponding pK*_a_* shifts, calculated from equation 1 and the experimental data of Din *et al.* ^37^

**Table 2:** Lipid A phosphate pK*_a_* shifts and theoretical values in the bilayer, as determined by free energy simulations.

| Bilayer | Charge state | $\Delta\Delta G$ | $\Delta pK_a$ | $pK_a$, theoretical |
| --- | --- | --- | --- | --- |
| Protonated | -4 → -5 | $6.8 \pm 0.3$ | $4.8 \pm 0.2$ | $13.4 \pm 0.2$ |
| | -5 → -6 | $7.3 \pm 0.3$ | $5.2 \pm 0.2$ | $15.9 \pm 0.2$ |
| Deprotonated | -4 → -5 | $2.7 \pm 0.6$ | $1.9 \pm 0.4$ | $10.5 \pm 0.4$ |
| | -5 → -6 | $1.0 \pm 0.4$ | $0.7 \pm 0.3$ | $11.5 \pm 0.3$ |

In all cases, the pK*_a_* shifts were positive, indicating an upward shift compared to the pK*_a_* in solution. The resultant theoretical pK*_a_* values were calculated as 10.5 and 11.5 for the first and second deprotonations when simulated in the deprotonated bilayer, and 13.4 and 15.9 when in the protonated bilayer. It should be noted that even a conservative reference pK*_a_* of 6.0, corresponding roughly to the pK*_a_* of glucose phosphate in water, ^63, 64^ would still result in lipid A pK*_a_* values for the first deprotonation well above physiological pH – 10.8 in the protonated bilayer and 7.9 in the deprotonated bilayer.

### NMR validates protonated LPS charge state

Given the large differences in simulated membrane properties between different LPS protonation states, experimental validation of the predicted pK*_a_* values is critical for informing force field paramaterizations. ^31^P NMR spectroscopy is sensitive to phosphate protonation state and the pH-dependence of chemical shifts can be used to determine pK*_a_* values.^65^ Magic angle spinning (MAS) ^31^P-NMR spectra for Re LPS show similar chemical shifts at pH 5.0, 7.0, and 9.0 (Figure 9) with peak positions around 1.3–1.4 ppm, demonstrating that no significant protonation state changes occur in this pH range. The spectra at pH 2.0 and 10.5, however, are shifted relative to the spectra at these intermediate pH values, with peak locations at −0.2 and 2.7 ppm, respectively. These chemical shift perturbations indicate changes in chemical environment for the phosphate groups at these extreme pHs, with pK_1_ between 2.0 and 5.0, and pK_2_ between 9.0 and 10.5 (highlighted regions in Figure 9B).

**Figure 9:**
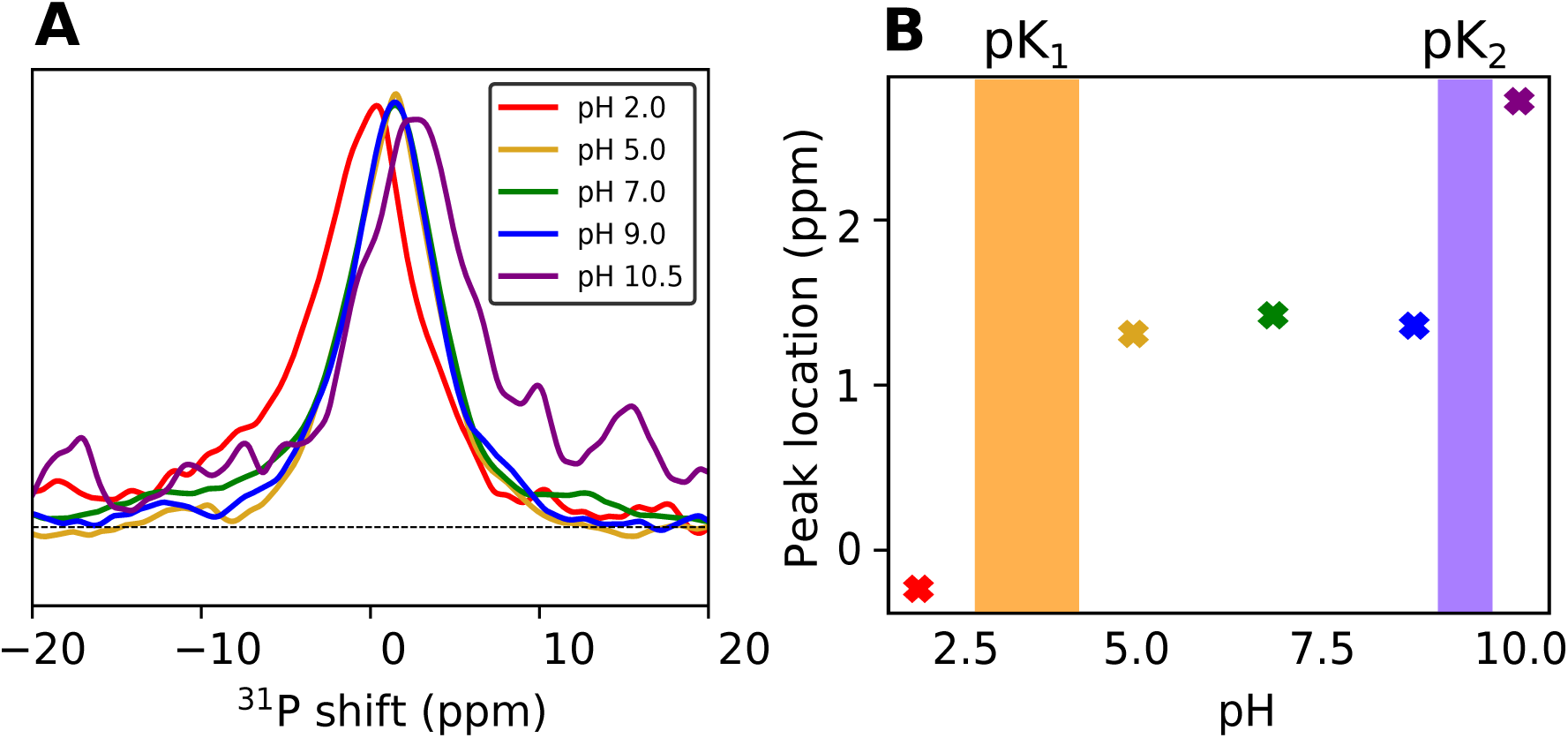
^31^P NMR pH titration of LPS. (A) ^31^P MAS spectra of LPS as a function of pH. Spectra of samples at five pH values are overlaid. (B) ^31^P peak position (in ppm) plotted as a function of sample pH. The two shaded regions indicate the putative approximate values for pK_1_ and pK_2_.

We propose pK_1_ represents the first ionization of the phosphate groups, from neutral to anionic, while pK_2_ represents the second ionization. The alternative, that the lipid A phosphate groups are deprotonated and have pK*_a_* values significantly lower than 5.0, is unlikely given that pK_1_ and pK_2_ of glucose phosphate are 1.0 and 6.1.^63, 64^ Furthermore, our interpretation of pK_1_ and pK_2_ is consistent with the known protonation state of Re LPS in solution as physiological pH, ^37^ and in good agreement with the predicted pK*_a_ ∼*10.5 from free energy simulations, while conventional simulations with singly-deprotonated LPS provide the best match to experimental structural data.

## Discussion

Discrepancies exist between the commonly-used LPS force fields, both in the lipid A phosphate group charges and in the bilayer response to the inclusion of different ions. The CHARMM LPS force field assigns a −2 net charge to each phosphate group ^25^ and displays significant compaction upon monovalent cation inclusion compared to divalent cations. ^34^ The GLYCAM LPS^23^ force field, on the other hand, models the phosphate groups as protonated (−1 charge) and correctly recovers the experimentally-observed condensing effect of divalent cations. In addition to the charge discrepancy, these force fields utilize different ion and water models as well.

In this work, we sought to understand the effects that altering the lipid A phosphate charges and including different ion parameterizations have on the structural characteristics of LPS simulations. Regardless of the phosphate charge utilized, use of the default CHARMM ion parameters^38^ with the standard sodium NBFIX terms^42^ resulted in smaller lipid areas with Na^+^ than Ca^2+^, in agreement with the simulation results of Kim *et al.* ^34^ This indicates that the differences in LPS response to ions between force fields cannot be attributed to the differing charge state alone, and that the ion or water parameters themselves are responsible. Indeed, implementation of the sodium/potassium CUFIX^39^ was sufficient in most systems to recover the expected trend of increased area and fluidity with monovalent cations.

Distinct differences were also observed between simulations with charges of −1 and −2 per lipid A phosphate group. Inter-lipid A hydrogen bonding was greatly increased for unmodified LPS systems when the protonated lipid A phosphates were utilized; this effect is likely a result of the additional hydrogen bond donors present in these systems. Furthermore, protonation of the phosphate groups had a condensing effect (Figures 5 and S4), especially striking in the LPS chemotypes. This decreased area per lipid resulted in more ordered, tightly packed lipid tails; the area per tail observed was 24.5–26.5 Å^2^ for Re LPS with protonated phosphate groups, compared to 26.5–30.5 Å^2^ for fully deprotonated Re LPS. These smaller lipid tail areas are in better agreement with measurements by Snyder *et al.*, which places the upper limit for the area per lipid tail at 26 Å^2^ for liquid-crystalline LPS.^20^ Though no experimental per lipid areas for the Rc or Re chemotypes could be found, the per lipid areas of 169 Å^2^ and 184 Å^2^ for protonated Rc LPS with Ca^2+^ and CUFIX Na^+^ are more consistent with the experimentally determined areas of 168.6 *±* 1.4 Å^2^ for the Ra chemotype with Ca^2+^ and 207.8 *±* 4.9 Å^2^ with Na^+^,^66^ compared to 179 and 175 Å^2^ in the corresponding deprotonated simulations. We note that, since the Ra chemotype contains more core sugars, the areas per lipid for the Rc chemotype are likely smaller than the experimental areas cited above. Overall, simulations demonstrate that models with protonated lipid A form more compact bilayers, which may represent a better match to the liquid-crystalline phase the outer membrane is believed to adopt physiologically. ^5,67^

Determination of the physiologically-relevant lipid A phosphate charge representation is a crucial problem that simulations need to address, especially in light of these differences observed between simulations with protonated or deprotonated lipid A phosphate groups. Here, free energy simulations were utilized to alchemically predict the pK*_a_* shift in the bilayer compared to solution, while ^31^P-NMR verified protonation state predictions. These results indicate that many LPS models, such as those utilized by the CHARMM-GUI and CG models based off it, incorrectly assign a charge of −2 to both lipid A phosphate groups which leads to significantly different bilayer properties. Furthermore, this charge assignment, especially when coupled with the strong nonbonded interactions of the CHARMM ions with oxygen, likely overestimates the affinity of cation binding. Since many AMPs are believed to competitively displace cations from the LPS core, ^68–70^ overly-stabilized cation interactions could have a significant effect on the outcome of such simulations.

Based on these results, we recommend that current LPS force field parameterizations be updated to correct for these charge inaccuracies, and offer the following phosphate charge guidelines: (1) At or near physiological pH, each LPS lipid A phosphate group should carry a charge of −1. (2) Phosphate groups adorned with positively-charged chemical modifications (AAB, PEtN, etc) should also carry a charge of −1. (3) Phosphate groups in the core, such as heptose-5 phosphate in the Re chemotype, are currently represented by a default charge of −2; future work is needed to definitively determine the appropriate protonation states of these groups. Furthermore, based on simulations with different ion parameterizations, we recommend use of the Na^+^/K^+^ CUFIX^39^ ion parameter sets when utilizing the CHARMM LPS force field for best agreement with experimental data. Finally, recent studies of phospholipids have revealed an overestimation of head group charge-charge interactions, which can lead to significant differences in bilayer properties; ^71^ while this study examined only the ion–phosphate interaction, LPS simulations may suffer from similar overestimation, and refinement of both intra-lipid and lipid–ion nonbonded parameters represent promising approaches for further model improvement. Additional calcium non-bonded refinement has occurred since the start of this work, ^72^ which may provide better agreement with experiment. Altogether, these results highlight inconsistencies in the current LPS models, while offering guidelines for choosing appropriate models to better reproduce experimental LPS results.

## Acknowledgement

The authors thank Prof. James Gumbart for helpful and constructive discussions concerning this work. Research reported in this publication was supported by the National Institute of General Medical Sciences of the National Institutes of Health (grant 1R35GM119647). The content is solely the responsibility of the authors and does not necessarily represent the official views of the National Institutes of Health. This work used the Extreme Science and Engineering Discovery Environment (XSEDE),^73^ Comet at the San Diego Supercomputer Center at UC San Diego through allocation TG-MCB140081.

## Supporting Information Available

Supplementary data analysis, tables, figures, and atomic partial charges for protonated lipid A phosphate groups.

---

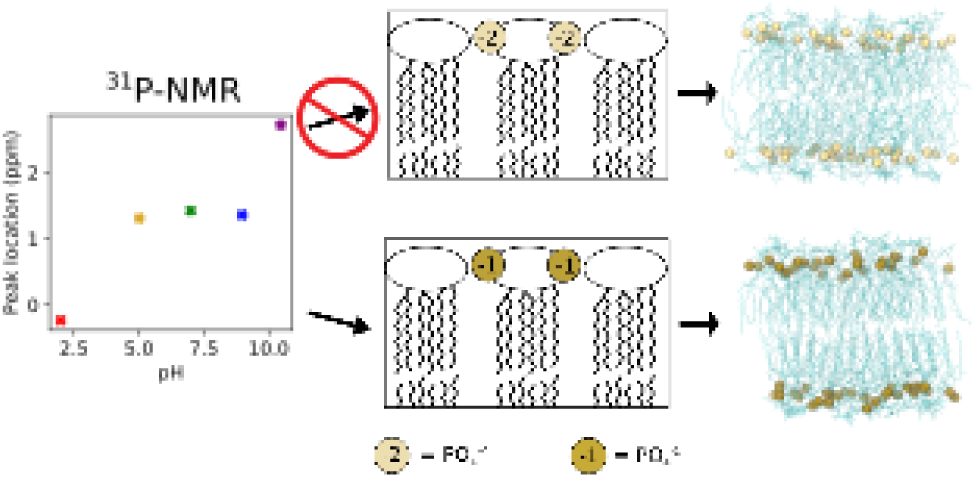
For Table of Contents Only.

