## Supporting Information for "Lipopolysaccharide Simulations are Sensitive to Phosphate Charge and Ion Parameterization"

**Content:**

14 pages of supplementary data
Sections S1 and S2 of supplementary text
Figures S1 to S7
Tables S1 to S6
References for SI reference citations

*

### S1 Simulations of deprotonated LPS with excess ions

To ensure that the differences observed between deprotonated and protonated LPS simulation results were not an artifact of the decreased number of counterions in the simulation box, four additional Re LPS simulations were performed. These simulations utilized the reduced phosphate charges, with the standard CHARMM $Na^+$, $K^+$, $Ca^{2+}$, or $Mg^{2+}$ ions. However, instead of adding counterions only to neutralize the system, an excess number of ions was included such that the number of cations in the simulation box was the same as the number used in the simulations with deprotonated lipid A phosphate groups: 192 or 96 for monovalent and divalent cations, respectively, instead of the 128 or 64 needed to neutralize. Additionally, 64 $Cl^-$ were added to the solvent to neutralize the simulation box. These four systems were simulated and analyzed as described in Materials and Methods.

No statistically significant differences were observed in the structural characteristics between systems when excess cations were included; the area per lipid, hydrophobic thickness, and inter-lipid A hydrogen bonding (Figure S1) were similar despite the presence of additional cations. Density profiles of the cation number density along the bilayer normal (Figure S2) reveal that the number of ions in the LPS core is similar in simulations with and without excess cations. Simulations with excess cations, however, display an increased density of cations in the bulk solvent. These results demonstrate that cations saturate the core in roughly the number needed to neutralize the anionic charges, while excess cations remain in the bulk solvent; this is likely why no significant differences were observed between simulations with differing number of cations present.

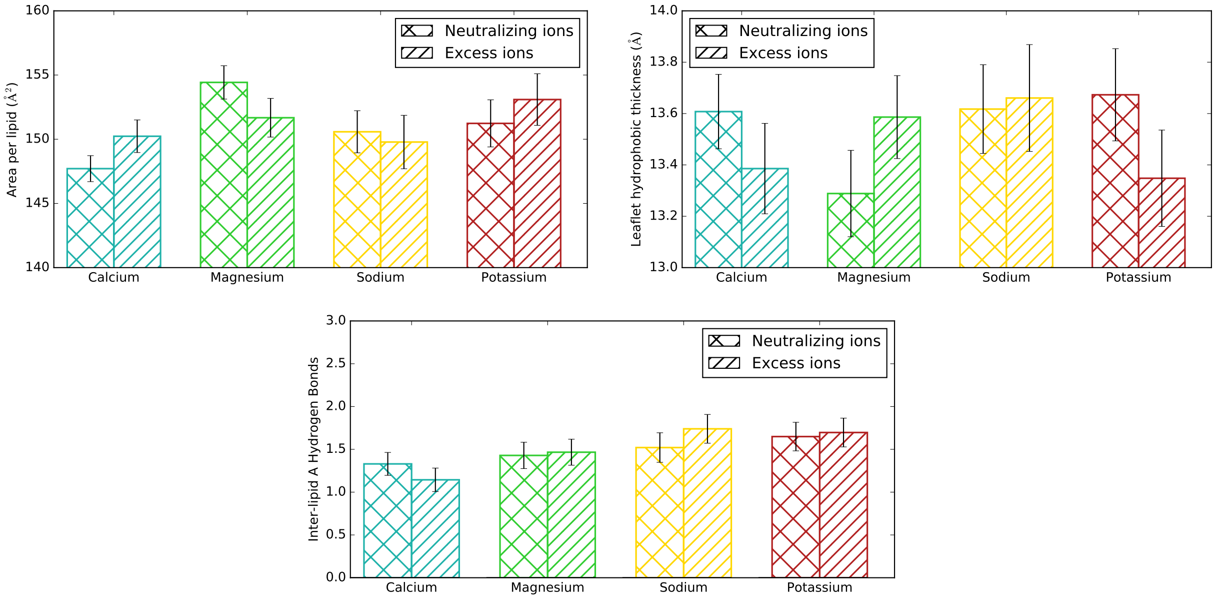


Figure S1: Area per lipid, hydrophobic thickness, and inter-lipid A hydrogen bonding for four Re LPS simulations with protonated lipid A phosphate groups. Addition of excess cations in the simulation box does not significantly affect the bilayer properties in these systems.

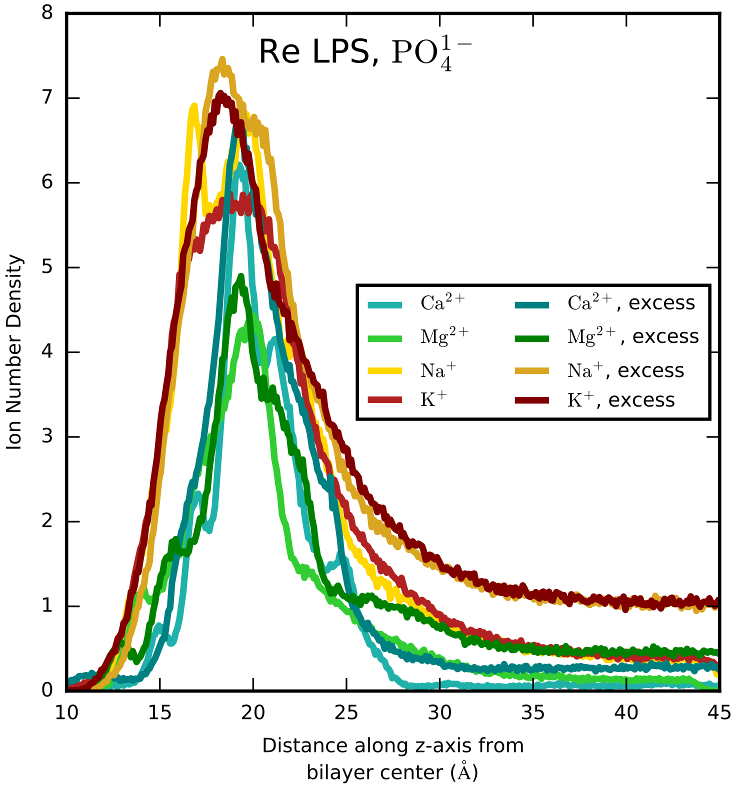


Figure S2: Ion number density along the bilayer normal for four Re LPS simulations with protonated lipid A phosphate groups, with and without excess cations in the simulation box. In all simulations, the LPS core contains roughly the number of ions needed to neutralize it; excess cations remain predominantly in the bulk solvent.

Table S1: Atom names, types, and charges for lipid A 1-P and 4'-P. Partial charges for $–PO_4^{2-}$ are from the parameter set of Wu *et al.* [1].

| Atom Name | Type, $–PO_4^{2-}$ | Charge, $–PO_4^{2-}$ | Type, $–PO_4^{-}$ | Charge, $–PO_4^{-}$ |
|---|---|---|---|---|
| CA1 | CC3162 | 0.11 | CC3162 | 0.21 |
| HA1 | HCA1 | 0.09 | HCA1 | 0.09 |
| OA1 | OC30P | -0.40 | OC30P | -0.62 |
| PA | PC | 1.10 | PC | 1.50 |
| OPA2 | OC2DP | -0.90 | OC2DP | -0.82 |
| OPA3 | OC2DP | -0.90 | OC312 | -0.67 |
| HPA3 | — | — | HCP1 | 0.33 |
| OPA4 | OC2DP | -0.90 | OC2DP | -0.82 |
| CB4 | CC3161 | -0.09 | CC3161 | -0.09 |
| HB4 | HCA1 | 0.09 | HCA1 | 0.19 |
| OB4 | OC30P | -0.40 | OC30P | -0.62 |
| PB | PC | 1.10 | PC | 1.50 |
| OPB2 | OC2DP | -0.90 | OC2DP | -0.82 |
| OPB3 | OC2DP | -0.90 | OC312 | -0.67 |
| HPB3 | — | — | HCP1 | 0.33 |
| OPB4 | OC2DP | -0.90 | OC2DP | -0.82 |

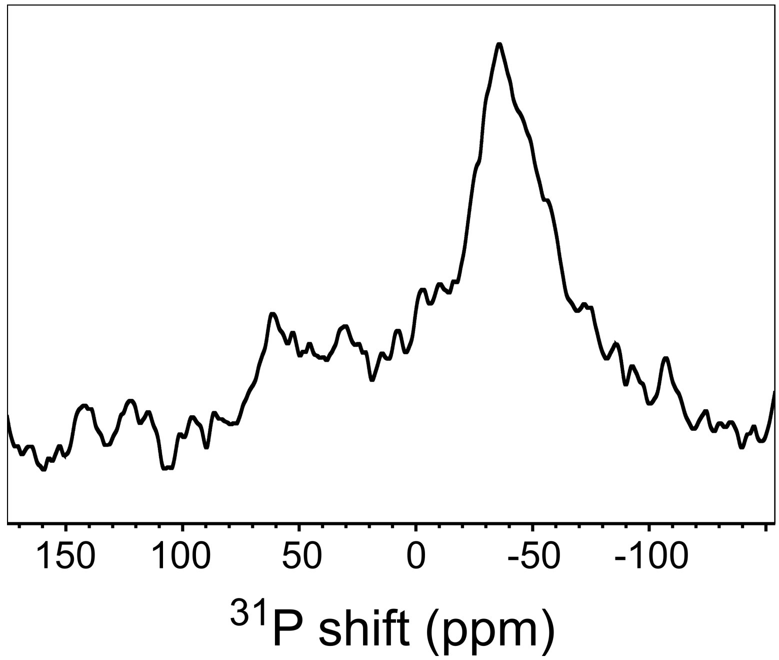


Figure S3: Static $^{31}$P-NMR spectrum at pH 7.0. The sample was studied in the hydrated state at 32 °C.

Table S2: Bilayer properties from simulations of LPS with $–PO_4^{2-}$ and different cation types. Error represents standard error of the mean.

| | | Are per lipid (Å$^2$) | Hydrophobic thickness (Å) | Inter-lipid hydrogen bonds |
|---|---|---|---|---|
| Re LPS | $Ca^{2+}$ | 163.7 ± 0.4 | 12.8 ± 0.1 | 0.59 ± 0.02 |
| | $Ca^{2+}$, NBFIX | 159.2 ± 0.5 | 13.2 ± 0.1 | 0.68 ± 0.12 |
| | $K^+$ | 167.5 ± 0.5 | 12.6 ± 0.1 | 0.56 ± 0.03 |
| | $K^+$, CUFIX | 181.9 ± 1.0 | 11.9 ± 0.1 | 0.36 ± 0.01 |
| | $Mg^{2+}$ | 169.8 ± 1.2 | 12.4 ± 0.1 | 0.61 ± 0.01 |
| | $Na^+$ | 153.8 ± 0.4 | 13.4 ± 0.1 | 0.89 ± 0.02 |
| | $Na^+$, CUFIX | 165.9 ± 0.8 | 12.7 ± 0.1 | 0.56 ± 0.03 |
| Rc LPS | $Ca^{2+}$ | 178.9 ± 0.2 | 11.9 ± 0.1 | 0.50 ± 0.02 |
| | $Ca^{2+}$, NBFIX | 176.5 ± 0.4 | 12.1 ± 0.1 | 0.45 ± 0.01 |
| | $K^+$ | 185.5 ± 0.7 | 11.6 ± 0.1 | 0.43 ± 0.02 |
| | $K^+$, CUFIX | 189.7 ± 0.8 | 11.5 ± 0.1 | 0.33 ± 0.01 |
| | $Mg^{2+}$ | 185.3 ± 0.4 | 11.9 ± 0.1 | 0.58 ± 0.02 |
| | $Na^+$ | 167.1 ± 0.3 | 12.3 ± 0.1 | 0.71 ± 0.02 |
| | $Na^+$, CUFIX | 175.3 ± 0.5 | 12.2 ± 0.1 | 0.50 ± 0.03 |
| Re mLPS | $Ca^{2+}$ | 181.6 ± 1.1 | 13.6 ± 0.1 | 1.66 ± 0.05 |
| | $Ca^{2+}$, NBFIX | 178.8 ± 0.3 | 13.7 ± 0.1 | 2.39 ± 0.09 |
| | $K^+$ | 179.0 ± 0.3 | 13.9 ± 0.1 | 2.75 ± 0.03 |
| | $K^+$, CUFIX | 178.0 ± 0.8 | 13.9 ± 0.1 | 2.17 ± 0.03 |
| | $Mg^{2+}$ | 182.1 ± 0.6 | 13.4 ± 0.1 | 2.57 ± 0.04 |
| | $Na^+$ | 176.2 ± 0.2 | 13.9 ± 0.1 | 2.28 ± 0.04 |
| | $Na^+$, CUFIX | 175.3 ± 0.4 | 14.0 ± 0.1 | 1.83 ± 0.05 |
| Rc mLPS | $Ca^{2+}$ | 186.1 ± 0.1 | 12.9 ± 0.1 | 2.25 ± 0.01 |
| | $Ca^{2+}$, NBFIX | 183.3 ± 0.6 | 13.4 ± 0.1 | 2.45 ± 0.04 |
| | $K^+$ | 180.4 ± 0.3 | 13.7 ± 0.1 | 2.75 ± 0.05 |
| | $K^+$, CUFIX | 192.1 ± 0.7 | 13.1 ± 0.1 | 1.88 ± 0.03 |
| | $Mg^{2+}$ | 191.4 ± 0.3 | 12.9 ± 0.1 | 2.49 ± 0.04 |
| | $Na^+$ | 177.1 ± 0.3 | 13.8 ± 0.1 | 2.31 ± 0.02 |
| | $Na^+$, CUFIX | 188.0 ± 0.4 | 13.4 ± 0.1 | 2.01 ± 0.02 |

Table S3: Bilayer properties from simulations of LPS with $–PO_4^-$ and different cation types. Error represents standard error of the mean.

| | | Are per lipid (Å$^2$) | Hydrophobic thickness (Å) | Inter-lipid hydrogen bonds |
|---|---|---|---|---|
| Re LPS | $Ca^{2+}$ | 147.7 ± 0.2 | 13.6 ± 0.1 | 1.33 ± 0.03 |
| | NBFIX $Ca^{2+}$ | 156.5 ± 0.6 | 13.2 ± 0.1 | 1.54 ± 0.02 |
| | $K^+$ | 151.2 ± 0.4 | 13.7 ± 0.1 | 1.65 ± 0.03 |
| | CUFIX $K^+$ | 159.9 ± 0.4 | 13.0 ± 0.1 | 1.54 ± 0.03 |
| | $Mg^{2+}$ | 154.4 ± 0.2 | 13.3 ± 0.1 | 1.43 ± 0.03 |
| | $Na^+$ | 150.6 ± 0.4 | 13.6 ± 0.1 | 1.52 ± 0.04 |
| | CUFIX $Na^+$ | 161.8 ± 0.4 | 12.8 ± 0.1 | 1.37 ± 0.05 |
| Rc LPS | $Ca^{2+}$ | 169.4 ± 0.3 | 12.5 ± 0.1 | 1.13 ± 0.02 |
| | NBFIX $Ca^{2+}$ | 173.0 ± 0.5 | 12.0 ± 0.1 | 1.25 ± 0.04 |
| | $K^+$ | 170.1 ± 1.5 | 12.5 ± 0.1 | 1.23 ± 0.03 |
| | CUFIX $K^+$ | 180.8 ± 1.1 | 12.0 ± 0.1 | 1.04 ± 0.05 |
| | $Mg^{2+}$ | 174.1 ± 1.3 | 12.2 ± 0.1 | 1.14 ± 0.03 |
| | $Na^+$ | 162.4 ± 0.5 | 12.8 ± 0.1 | 1.32 ± 0.6 |
| | CUFIX $Na^+$ | 183.7 ± 1.5 | 11.8 ± 0.1 | 0.87 ± 0.02 |
| Re mLPS | $Ca^{2+}$ | 174.4 ± 0.7 | 14.0 ± 0.1 | 2.04 ± 0.02 |
| | NBFIX $Ca^{2+}$ | 176.9 ± 0.3 | 13.9 ± 0.1 | 2.20 ± 0.03 |
| | $K^+$ | 172.2 ± 0.8 | 14.3 ± 0.1 | 2.67 ± 0.04 |
| | CUFIX $K^+$ | 175.7 ± 0.5 | 14.1 ± 0.1 | 2.14 ± 0.03 |
| | $Mg^{2+}$ | 172.0 ± 0.3 | 14.3 ± 0.1 | 2.64 ± 0.03 |
| | $Na^+$ | 173.9 ± 0.7 | 14.0 ± 0.1 | 2.48 ± 0.02 |
| | CUFIX $Na^+$ | 174.8 ± 0.2 | 14.1 ± 0.1 | 2.10 ± 0.03 |
| Rc mLPS | $Ca^{2+}$ | 177.9 ± 0.1 | 13.9 ± 0.1 | 1.91 ± 0.02 |
| | NBFIX $Ca^{2+}$ | 183.0 ± 0.5 | 13.4 ± 0.1 | 2.21 ± 0.02 |
| | $K^+$ | 179.7 ± 0.8 | 13.5 ± 0.1 | 2.50 ± 0.04 |
| | CUFIX $K^+$ | 185.3 ± 0.7 | 13.7 ± 0.1 | 1.86 ± 0.03 |
| | $Mg^{2+}$ | 184.5 ± 0.3 | 13.7 ± 0.1 | 2.17 ± 0.04 |
| | $Na^+$ | 179.2 ± 0.7 | 13.3 ± 0.1 | 2.24 ± 0.07 |
| | CUFIX $Na^+$ | 185.3 ± 0.5 | 13.5 ± 0.1 | 1.81 ± 0.04 |

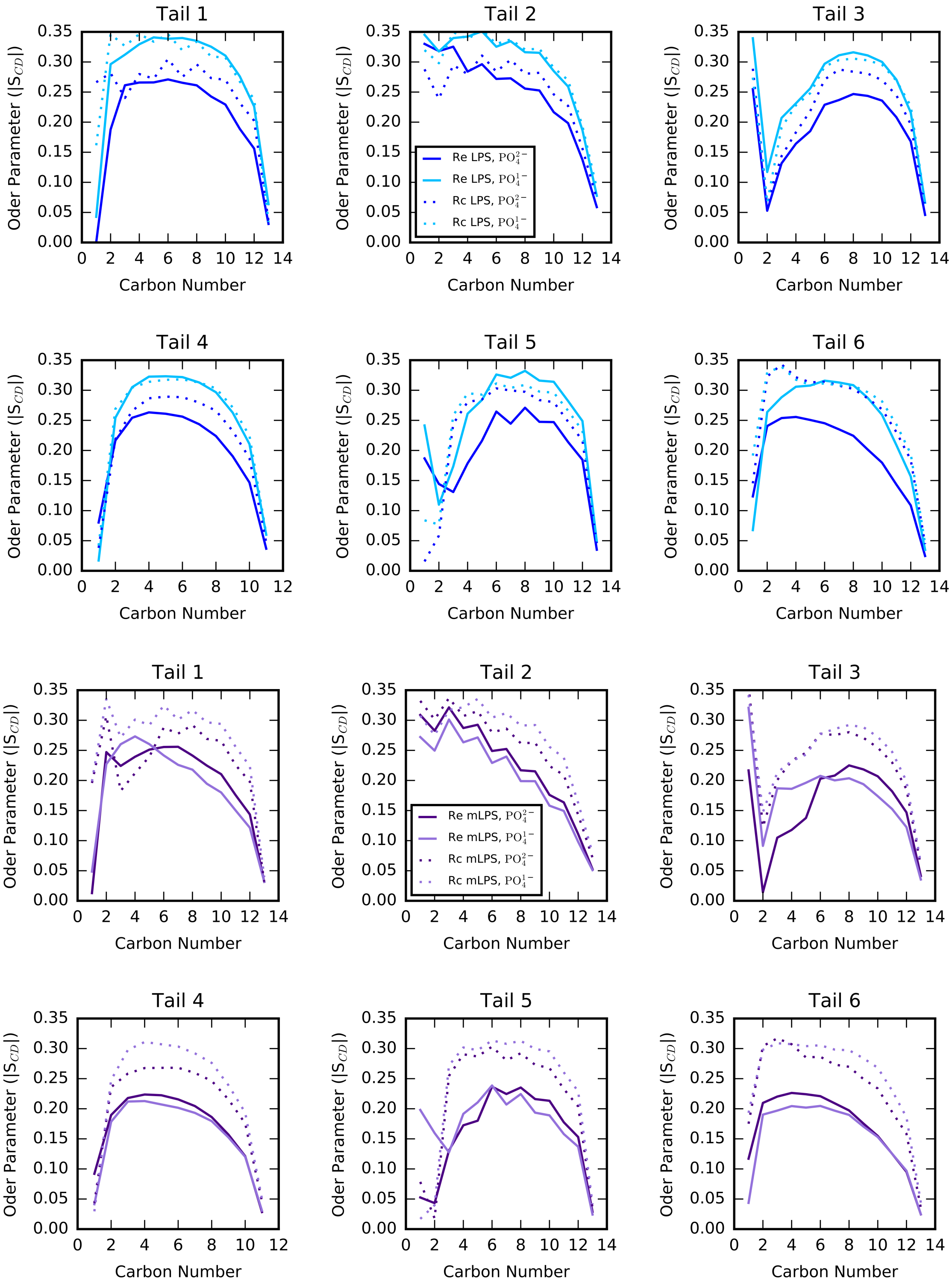


Figure S4: Lipid tail order parameters (|S$_{cd}$|) for all six lipid tails. Data for LPS systems are shown in shades of blue, while data for mLPS systems are in shades of purple. Lighter colors denote systems with –$PO_4^-$.

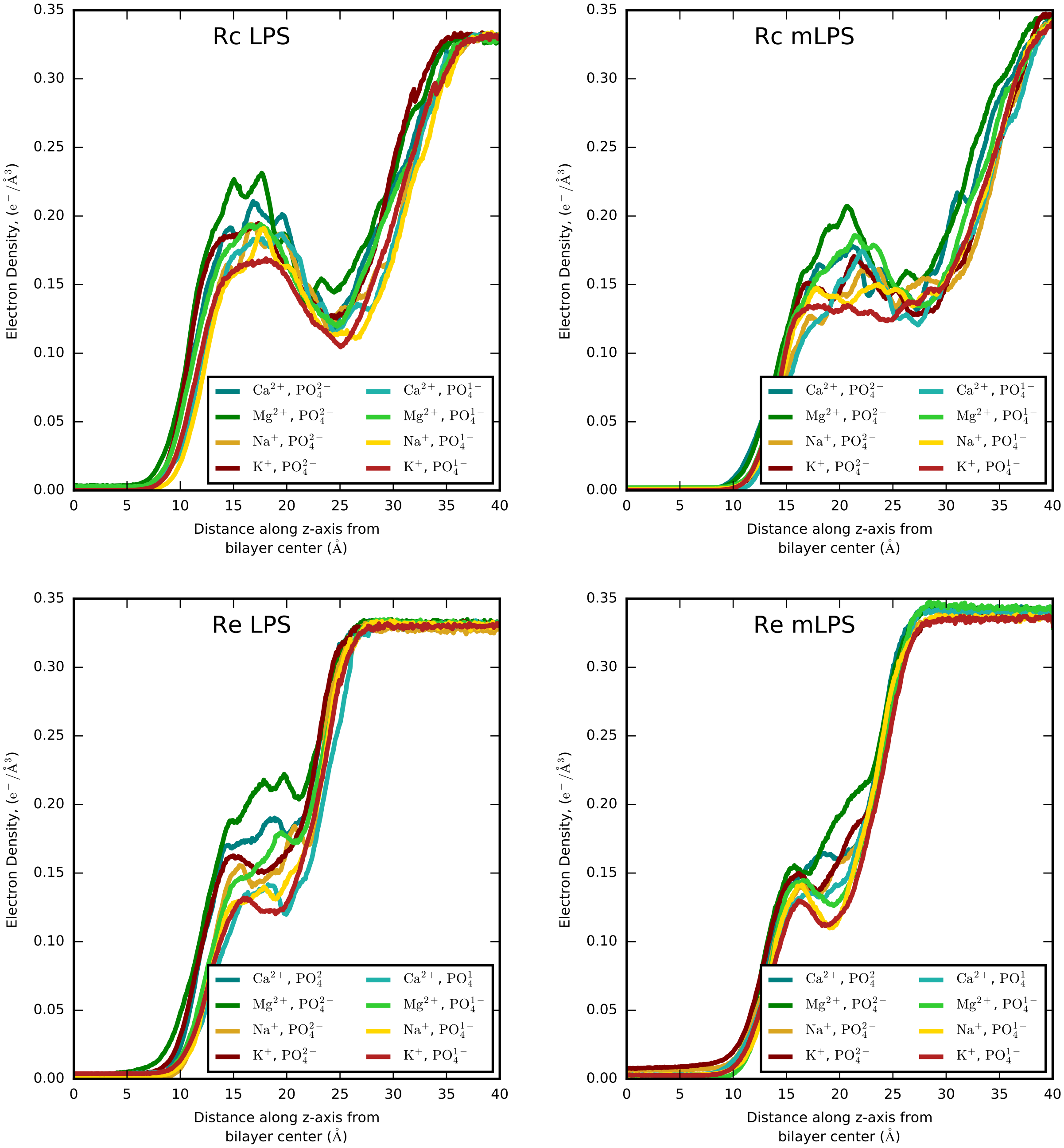


Figure S5: Water electron density for all four chemotypes simulated. Simulations with reduced phosphate charges (lighter shades) display less saturated LPS core compared to simulations with deprotonated lipid A phosphate groups.

Table S4: Average water electron density ($e^-/\text{Å}^3$) in the lipid A phosphate group region for all Re simulations. Utilization of reduced phosphate charges leads to decreased water density within the LPS core, regardless of LPS or ion type.

| | Re LPS | | Re mLPS | |
|---|---|---|---|---|
| | $-PO_4^{-2}$ | $-PO_4^-$ | $-PO_4^{-2}$ | $-PO_4^-$ |
| $Ca^{2+}$ | 0.176 ± 0.007 | 0.147 ± 0.008 | 0.162 ± 0.007 | 0.133 ± 0.006 |
| NBFIX $Ca^{2+}$ | 0.179 ± 0.007 | 0.163 ± 0.007 | 0.156 ± 0.007 | 0.137 ± 0.008 |
| $K^+$ | 0.161 ± 0.008 | 0.122 ± 0.007 | 0.138 ± 0.007 | 0.114 ± 0.006 |
| CUFIX $K^+$ | 0.186 ± 0.007 | 0.145 ± 0.007 | 0.132 ± 0.005 | 0.131 ± 0.007 |
| $Mg^{2+}$ | 0.208 ± 0.009 | 0.150 ± 0.007 | 0.166 ± 0.007 | 0.127 ± 0.006 |
| $Na^+$ | 0.144 ± 0.007 | 0.134 ± 0.006 | 0.135 ± 0.007 | 0.116 ± 0.006 |
| CUFIX $Na^+$ | 0.170 ± 0.007 | 0.162 ± 0.007 | 0.131 ± 0.008 | 0.130 ± 0.008 |

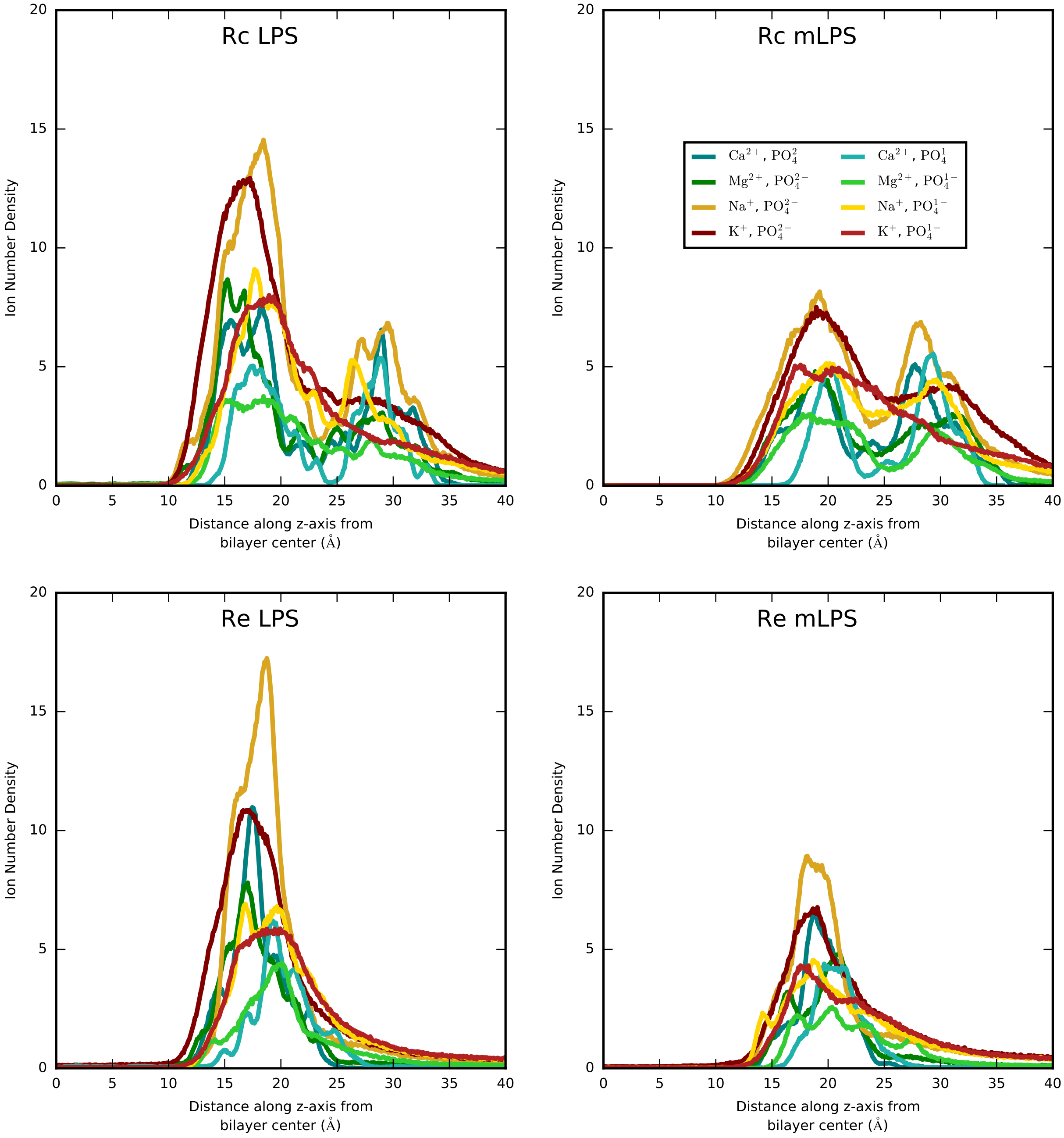


Figure S6: Ion number density along the bilayer normal for all four chemotypes simulated. In all simulations, the ions remain predominantly in the LPS core rather than in the bulk solvent.

Table S5: Average free energy for each deprotonation step. Error represents the standard error of the mean. All values are in kcal/mol.

| | Water | Protonated LPS bilayer | Deprotonated LPS bilayer |
|---|---|---|---|
| 1st deprotonation, $P_A$ first ($\Delta G_{1,P_A}$) | -99.0 ± 0.1 | -92.2 ± 0.2 | -87.0 ± 0.6 |
| 2nd deprotonation, $P_A$ first ($\Delta G_{2,P_A}$) | -126.7 ± 0.1 | -119.4 ± 0.2 | -129.9 ± 0.9 |
| 1st deprotonation, $P_B$ first ($\Delta G_{1,P_B}$) | -126.8 ± 0.2 | -119.0 ± 0.6 | -124.1 ± 0.6 |
| 2nd deprotonation, $P_B$ first ($\Delta G_{2,P_B}$) | -98.9 ± 0.1 | -99.3 ± 0.4 | -97.9 ± 0.4 |

Table S6: Lennard-Jones parameters for all atom pairs modified by NBFIX or CUFIX. The standard parameters are computed by the Lorentz-Berthelot combination rule and included for comparison. All changes affect $R_{min}$ only. Atom type OBL corresponds to the ester =O in the lipid tails, OC2D2 is carboxylate =O present in KDO in these simulations, and OC2DP is the phosphate =O of all phosphate groups. † Default CHARMM parameters of Beglov and Roux [2]. * Sodium NBFIX parameters of Venable *et al.* [3], which are included in the standard CHARMM ion parameter set. ‡ NBFIX calcium parameters of Roux and Rong as reported by Kim *et al.* [4]. ° CUFIX parameters of Yoo and Aksimentiev [5].

| Ion | Set Name | Atom 1 | Atom 2 | $R_{min}$ (Å) |
|---|---|---|---|---|
| Sodium | NBFIX (default)* | SOD | OBL | 3.13000 |
| | NBFIX (default)* | SOD | OC2D2 | 3.23000 |
| | CUFIX° | SOD | OC2D2 | 3.20075 |
| | NBFIX (default)* | SOD | OC2DP | 3.16000 |
| | CUFIX° | SOD | OC2D2 | 3.20075 |
| Calcium | NBFIX‡ | CAL | OC2D2 | 3.22500 |
| | Standard† | CAL | OC2D2 | 3.06700 |
| | NBFIX‡ | CAL | OC2DP | 3.30400 |
| | Standard† | CAL | OC2DP | 3.06700 |
| Potassium | CUFIX° | POT | OC2D2 | 3.54375 |
| | Standard† | POT | OC2D2 | 3.46375 |
| | CUFIX° | POT | OC2DP | 3.54375 |
| | Standard† | POT | OC2DP | 3.11075 |

#### S2 Comparison of 4x4 and 6x6 LPS systems

Results from simulations of 4x4 Rc LPS and mLPS systems are consistent with our previously-published 6x6 simulations [6]. Two additional 6x6 Rc LPS simulations for comparison- one with protonated (-1) lipid A phosphate and with deprotonated (-2) lipid A phosphate groups. Additionally, the Monte Carlo barostat was utilized in these new simulations. All other simulation conditions were the same; each simulation was performed for 3.0 µs with analysis over the last 1.0 µs as before. The areas per lipid for the two new simulations are shown in Figure S7 below, along with the corresponding 4x4 systems for comparison. We find there is no significant difference in the area per lipid, hydrophobic thickness, or tail order parameters. For example, the area per lipid for the protonated 4x4 LPS system is 173.0 ± 0.5 Å$^2$, compared to 173.2 ± 0.9 Å$^2$ for the 6x6 system with the MC barostat. The deprotonated systems are also statistically indistinguishable- 176.5 ± 0.4 Å$^2$ for the 4x4 system and 176.0 ± 0.6 Å$^2$ for the 6x6 system.

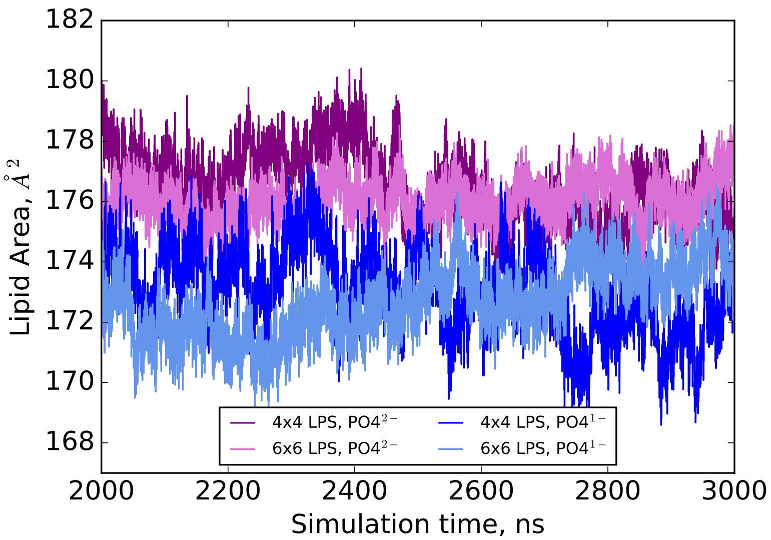


Figure S7: Comparison of 4x4 and 6x6 LPS systems with Berendsen or Monte Carlo barostat methods, respectively. Area per lipid is given over the last µs of simulations of Rc LPS with NBFIX calcium. Purple shades are simulations with $PO_4^{2-}$ while blue shades are simulations with $PO_4^{1-}$.
